# MNS induces antiviral protection and suppresses inflammation

**DOI:** 10.64898/2026.02.11.705318

**Authors:** Yang Zhao, Xingyu Chen, Yunfei Xie, Hanjie Liu, Boming Kang, Shuailong Zheng, Yuxiang Ren, Qian Wang, Fuping You, Haoxiang Qi

**Author notes:** These authors share the first authorship (Yang Zhao, Xingyu Chen, Yunfei Xie, Hanjie Liu). **Correspondence:** Haoxiang Qi.

## Abstract

**Background:** Identifying safe and broad-spectrum antiviral and anti-inflammatory agents remains an urgent need in infectious and inflammatory diseases. Here, we demonstrated that MNS (NSC170724), a small-molecule nitrovinyl benzodioxole, enhanced antiviral defense while limiting excessive inflammation.

**Methods:** The antiviral activity of MNS was evaluated in multiple cell lines and mouse infection models across DNA and RNA viruses. Virus-induced and LPS-induced inflammatory responses were assessed using RT-qPCR, ELISA and western blotting. Bulk RNA-seq and ATAC-seq were performed to define transcriptional and epigenetic mechanisms.

**Results:** MNS significantly suppressed viral infection in vitro and improved survival in four lethal viral infection models, accompanied by reduced viral loads and attenuated tissue injury. MNS also diminished virus-triggered and LPS-triggered inflammatory cytokine production in macrophages and multiple mouse organs, and protected mice from LPS-induced endotoxic lethality. Multi-omics profiling showed that MNS broadly repressed LPS-induced inflammatory transcriptional programs and reversed chromatin accessibility gains across promoters and transcription start sites. Joint analysis of RNA-seq and ATAC-seq data demonstrated consistent downregulation of pivotal inflammatory pathways, such as NF-κB, Toll-like receptor, and TNF signaling.

**Conclusions:** With potent activity against viral replication and inflammation in cellular and animal models, MNS emerges as a promising candidate for the treatment of viral infections and hyperinflammatory conditions.

## Introduction

Viral infections remain a major cause of morbidity and mortality worldwide, as illustrated by seasonal epidemics, emerging and re-emerging pathogens, and the ongoing threat of pandemic outbreaks [1–3]. In many settings, current antiviral therapies are limited by narrow specificity, the need for early administration, and the rapid emergence of viral resistance [4, 5]. Most approved agents directly target viral proteins or enzymes, which constrains their spectrum of activity and may fail when viruses accumulate escape mutations [6, 7]. At the same time, severe disease is often driven not only by uncontrolled viral replication but also by dysregulated host responses, including hyperinflammation, endothelial damage, and multiorgan failure [8–10]. This dual contribution of viral burden and host pathology underscores the need for strategies that both enhance antiviral defense and maintain immune homeostasis rather than simply blocking one viral target.

Host-directed therapies have therefore attracted growing interest as an alternative or complementary approach to classic direct-acting antivirals [11, 12]. By modulating cellular pathways that are essential for viral replication or innate immune defense, host-directed agents have the potential to provide broader coverage across different viruses and to retain activity despite viral evolution [13, 14]. Several experimental and clinical agents target pattern-recognition receptor signaling, interferon pathways, or downstream effector modules to boost antiviral immunity [15, 16]. However, many of these approaches focus on enhancing immune activation and may carry a risk of amplifying tissue-damaging inflammation if not tightly regulated [17, 18]. An ideal host-directed drug would combine antiviral efficacy with intrinsic anti-inflammatory properties, strengthening antiviral programs while preventing excessive cytokine production and tissue injury.

Macrophages sit at the center of this balance between protection and pathology [19]. As professional sentinels of the innate immune system, they sense viral components and bacterial products such as lipopolysaccharide (LPS) through Toll-like receptors, RIG-I–like receptors, and other pattern-recognition receptors [20–22]. Activation of these pathways rapidly induces transcription factors such as NF-κB and IRFs, leading to robust production of type I interferons, interleukins, chemokines, and other mediators that coordinate antiviral and inflammatory responses [23, 24]. While these programs are essential for early pathogen control, their sustained or excessive activation can drive systemic inflammation, vascular leakage, and organ damage in conditions such as viral sepsis, acute respiratory distress syndrome, and septic shock [25, 26]. Pro-inflammatory cytokines including IL-1β, IL-6, and TNF-α, as well as effector enzymes like COX-2 and iNOS, have been strongly linked to poor outcomes in hyperinflammatory states [27, 28]. The NLRP3 inflammasome further amplifies these responses by promoting IL-1β maturation and pyroptotic cell death [29, 30]. Drugs that can dampen these pathways without abolishing host defense are therefore of considerable therapeutic interest.

Recent researches have revealed that these innate immune responses are not governed solely by signaling cascades, but also by dynamic changes in the transcriptional and epigenetic landscape [31]. Inflammatory stimulation leads to rapid remodeling of chromatin accessibility at promoters and enhancers, recruitment of transcription factors, and deposition or removal of histone marks [32]. These changes shape the magnitude and kinetics of cytokine gene induction and can leave a form of “inflammatory memory,” contributing to tolerance or trained immunity depending on context [33, 34]. The transcriptional and chromatin dynamics of innate immunity can now be characterized genome-wide, enabled by techniques including bulk RNA sequencing (RNA-seq) and assay for transposase-accessible chromatin with sequencing (ATAC-seq) [35]. Studies using these tools have mapped how LPS and other stimuli reprogram macrophage gene networks, but how small-molecule host-directed drugs rewire chromatin accessibility and inflammatory transcriptional programs remains incompletely understood [36]. Dissecting these mechanisms is important for rationally designing agents that prevent harmful inflammation while preserving essential antimicrobial functions.

MNS (NSC170724) is a small-molecule nitrovinyl benzodioxole that was originally characterized as an orally available tyrosine kinase inhibitor and broad antiplatelet compound [37]. It inhibits kinases such as Src, Syk, and FAK, which participate in integrin signaling, immune receptor pathways, and cytoskeletal dynamics [38]. Recent preclinical studies further indicate that MNS inhibits both NLRP3 inflammasome activation and β1 integrin signaling, thereby mitigating NLRP3-driven inflammatory conditions such as experimental colitis and ischemia–reperfusion injury [39]. These findings suggest that MNS can modulate innate immune signaling and inflammatory effector pathways in vivo. However, its potential role in antiviral defense, as well as its impact on systemic inflammation induced by viral infection or endotoxemia, has not been systematically evaluated. Moreover, the transcriptional and epigenomic mechanisms by which MNS might exert coordinated antiviral and anti-inflammatory effects remain unclear.

Large-scale empirical screening to identify molecules with both antiviral and anti-inflammatory properties is labor-intensive and costly. Computational approaches that integrate biological knowledge and high-dimensional data offer an efficient way to prioritize candidate compounds.

We recently developed DeepAVC, a deep-learning framework that combines phenotype- and target-guided information to predict broad-spectrum antiviral compounds [40]. Using this platform, we identified MNS as a candidate host-directed antiviral agent. Its known ability to inhibit kinases involved in immune signaling and to restrain NLRP3 inflammasome activation made it an attractive molecule to test the hypothesis that a single small molecule could both enhance antiviral protection and limit excessive inflammation.

In the present study, we investigated the antiviral and anti-inflammatory activities of MNS using complementary in vitro and in vivo models. We first assessed whether MNS could inhibit replication of representative RNA and DNA viruses in different cell types, and whether it could improve survival and reduce viral burden and tissue injury in mouse models of lethal viral infection. We then examined its ability to attenuate virus-induced and LPS-induced inflammatory responses in macrophages and in a systemic LPS-induced sepsis model. Finally, to delineate the impact of MNS on LPS-induced inflammation, we performed bulk RNA-seq and ATAC-seq analyses, focusing on essential innate immune pathways including NF-κB, Toll-like receptor, and TNF signaling, and assessed associated changes in gene expression and chromatin accessibility. By integrating functional virology, inflammation models, and multi-omics profiling, we aimed to determine whether MNS acts as a host-protective agent that couples broad antiviral activity with epigenetically mediated suppression of hyperinflammatory responses, and to provide mechanistic insight into its mode of action.

## Result

### MNS Establishes a Host-Protective Antiviral State in vitro

Leveraging our previous established DeepAVC framework-a tool for precise, phenotype- and target-directed prediction of broad-spectrum antiviral agents [40]. We nominated MNS as a lead molecule and systematically examined its activity in cell-based assays. RAW 264.7 macrophages and bone-marrow–derived macrophages (BMDMs) were exposed to 12.5 or 25 μM MNS for 12 h, with DMSO-treated cells serving as controls, and were subsequently infected at an MOI of 0.1 with vesicular stomatitis virus (VSV), herpes simplex virus type 1 (HSV-1), mouse hepatitis virus (MHV) or encephalomyocarditis virus (EMCV) (Figure 1A-1B). THP-1 and HT1080 cells underwent the same treatment and infection regimen with VSV, HSV-1 or EMCV (Figure 1C-1D). RT-qPCR showed a clear concentration-dependent decline in viral mRNA in MNS-treated cultures relative to DMSO controls, with maximal inhibition at 25 μM. Consistently, plaque assays demonstrated markedly reduced infectious virus production in all examined cell types following MNS exposure, confirming robust antiviral activity in vitro. Notably, the magnitude of viral RNA reduction was more modest than the pronounced decrease in infectious virus production measured by plaque assays, suggesting that MNS primarily impairs the generation of infectious progeny rather than completely abolishing viral RNA synthesis.

**Figure 1.**
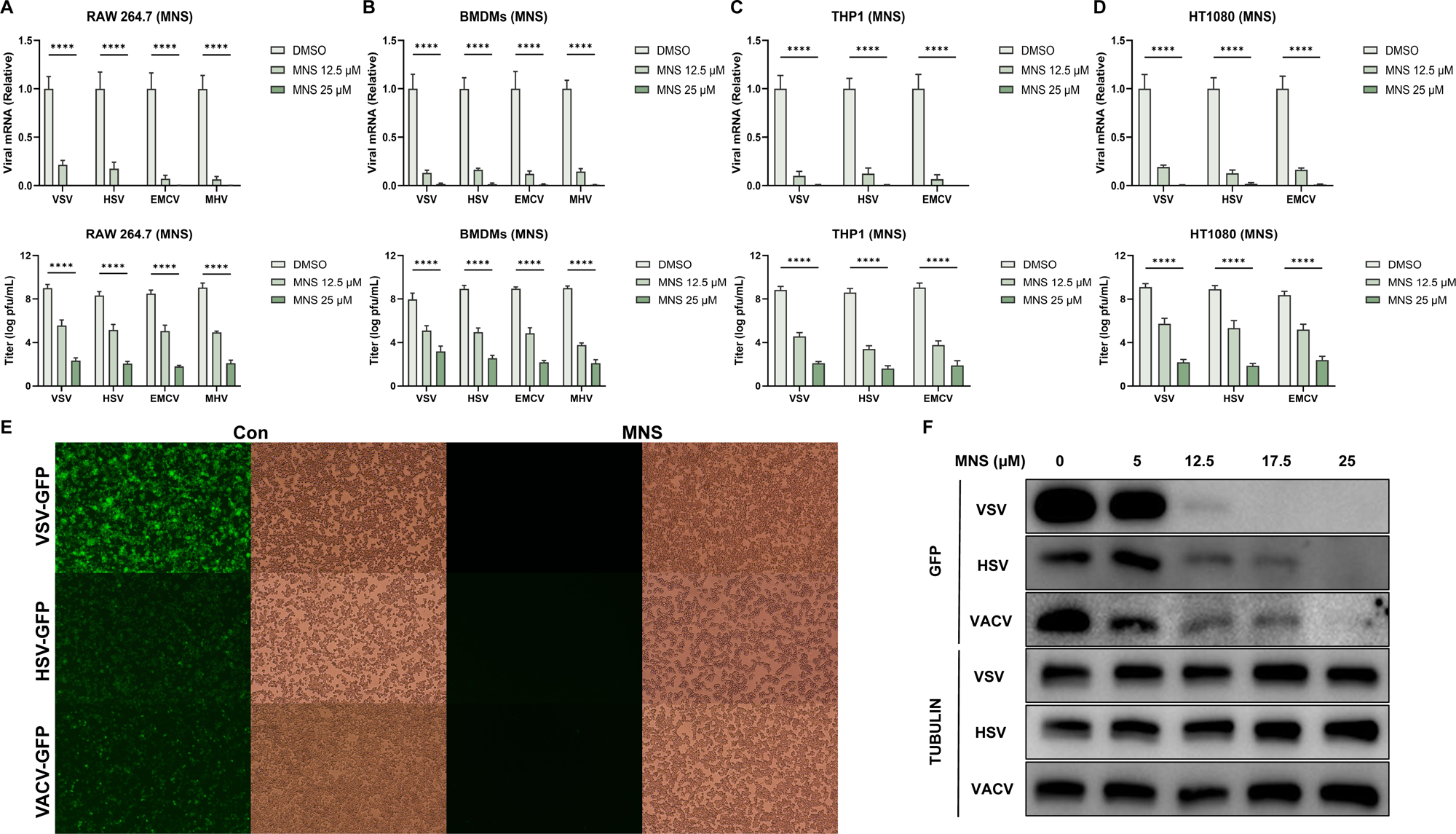
MNS Establishes a Host-Protective Antiviral State in vitro. (A-D) RT-qPCR quantification of viral RNA and plaque assay of viral titers in RAW264.7 cells (A), BMDMs (B), THP1 cells (C) and HT1080 cells (D) infected with viruses following the treatment of MNS. (E) Representative fluorescence/bright□field images of RAW 264.7 cells pretreated with DMSO or 25□µM MNS and infected with GFP□tagged VSV, HSV□1, or VACV (MOI□=□0.1, 12□h). (F) Dose□dependent reduction of viral GFP expression (VSV, HSV□1, VACV) by MNS in RAW 264.7 cells, analyzed by western blot (GFP/Tubulin) after infection (MOI□=□0.1, 12□h). Data: mean□±□SEM. N.S., p□>□0.05, *p□<□0.05, **p□<□0.01, ***p□<□0.001.

To characterize the spectrum of this effect, we next infected cells with GFP-reporter viruses, including DNA viruses (HSV-1 and vaccinia virus [VACV]) and an RNA virus (VSV). Fluorescence microscopy revealed pronounced loss of GFP signal after MNS treatment (Figure 1E), indicating efficient blockade of viral replication across distinct viral families. In line with this, western blot analysis of GFP in RAW 264.7 cells infected with HSV-1-GFP, VACV-GFP or VSV-GFP demonstrated a dose-responsive suppression of viral protein expression (Figure 1F). Antiviral activity was already evident at 5 μM MNS, and GFP signals were nearly undetectable at 25 μM, reflecting near-complete inhibition of replication.

Moreover, dose-response analyses revealed that MNS inhibited viral replication in a concentration-dependent manner in RAW 264.7 cells and HT1080 cells, with EC50 values in the low micromolar range across VSV and HSV-1 infection models. In contrast, MNS exhibited minimal cytotoxicity in the same cell types, with CC50 values exceeding the highest concentrations tested. As a result, MNS displayed favorable selectivity indices, indicating that its antiviral activity was not attributable to impaired cell viability (Supplementary Figure 1A-1B).

Collectively, these findings indicated that MNS induced a robust antiviral program that shields diverse host cell types from viral infection in vitro.

### MNS Establishes a Host-Protective Antiviral State in vivo

Building upon these promising in vitro results, we next investigated the in vivo protective efficacy of MNS. Following intraperitoneal infection with HSV-1, VSV, EMCV, or MHV, six-week-old C57BL/6J mice received intraperitoneal administration of either MNS or a vehicle control. Survival was recorded over a 7-day period. Across all four viral infection models, mice in the MNS-treated group exhibited a markedly higher survival rate than those in the control group. (Figure 2A-2D).

**Figure 2.**
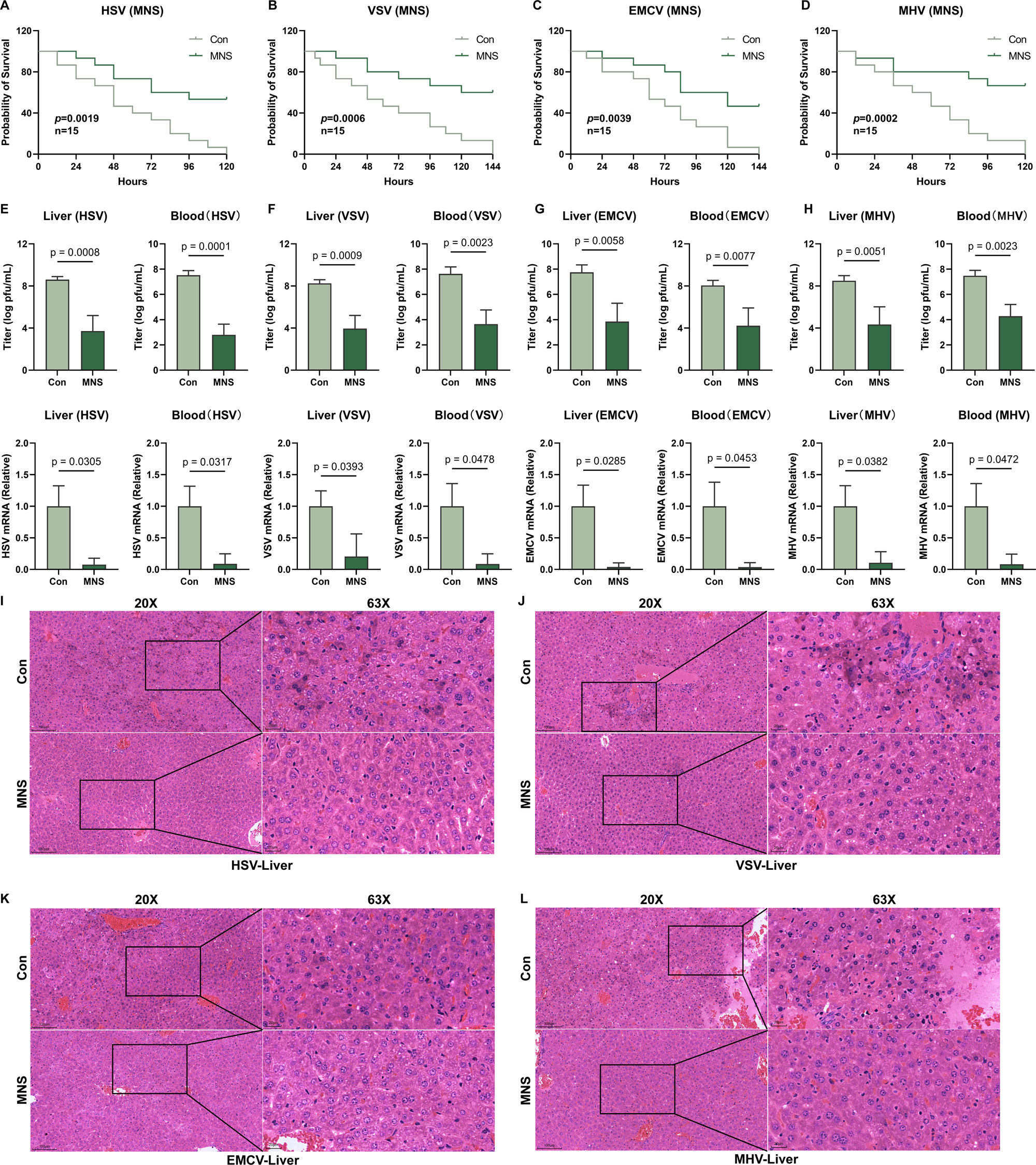
MNS Establishes a Host-Protective Antiviral State in vivo. (A□D) MNS conferred protection against lethal viral challenge in C57BL/6J mice. Kaplan□Meier survival plots show mice infected i.p. with HSV□1 (A), VSV (B), EMCV (C), or MHV (D) (n = 15 per group) and treated daily with vehicle or MNS (5 mg/kg). (E□H) Viral RNA (RT□qPCR) and viral titers (plaque assay) measured in blood and liver collected 24 h post□infection with HSV□1 (E), VSV (F), EMCV (G), or MHV (H). (I-L) Histopathology of liver sections from mice infected and treated as in (E□H). Low□ (20×) and high□magnification (63×, boxed areas) views are shown (scale bars: 100□µm and 20□µm). Data are mean□±□SEM; N.S., p□>□0.05; *p□<□0.05, **p□<□0.01, ***p□<□0.001.

To assess control of viral infection, we collected blood and liver tissues from a subset of mice at 24 hours post-infection. Plaque assays and RT-qPCR analysis consistently showed markedly reduced viral loads and viral mRNA levels in the MNS-treated groups across all tested viruses (Figure 2E-2H), confirming robust antiviral activity in vivo. Histological examination of liver sections further supported MNS-mediated protection, revealing attenuated inflammatory cell infiltration and tissue damage in MNS-treated mice infected with HSV-1, VSV, EMCV or MHV compared with vehicle controls (Figure 2I-2L).

Collectively, our results demonstrated that MNS-treatment induced a robust host-protective state characterized by broad-spectrum antiviral effects, effectively limiting viral infection and virus-induced tissue damage in vivo.

### MNS Suppressed Virus-induced and LPS-induced Inflammatory Responses

Having established the strong antiviral activity of MNS, we next examined whether MNS also reduced virus-induced inflammatory responses. In vitro, MNS treatment significantly decreased the induction of inflammatory cytokines following infection with VSV, HSV-1, EMCV or MHV in RAW264.7 cells, BMDMs and THP-1 cells. RT-qPCR analysis showed markedly reduced *IL1B* mRNA levels in cells treated with 12.5 μM or 25 μM MNS compared with DMSO controls. Consistently, ELISA measurements demonstrated a dose-dependent reduction in secreted IL-1β protein levels across all four viral infections (Figure 3A-3C).

**Figure 3.**
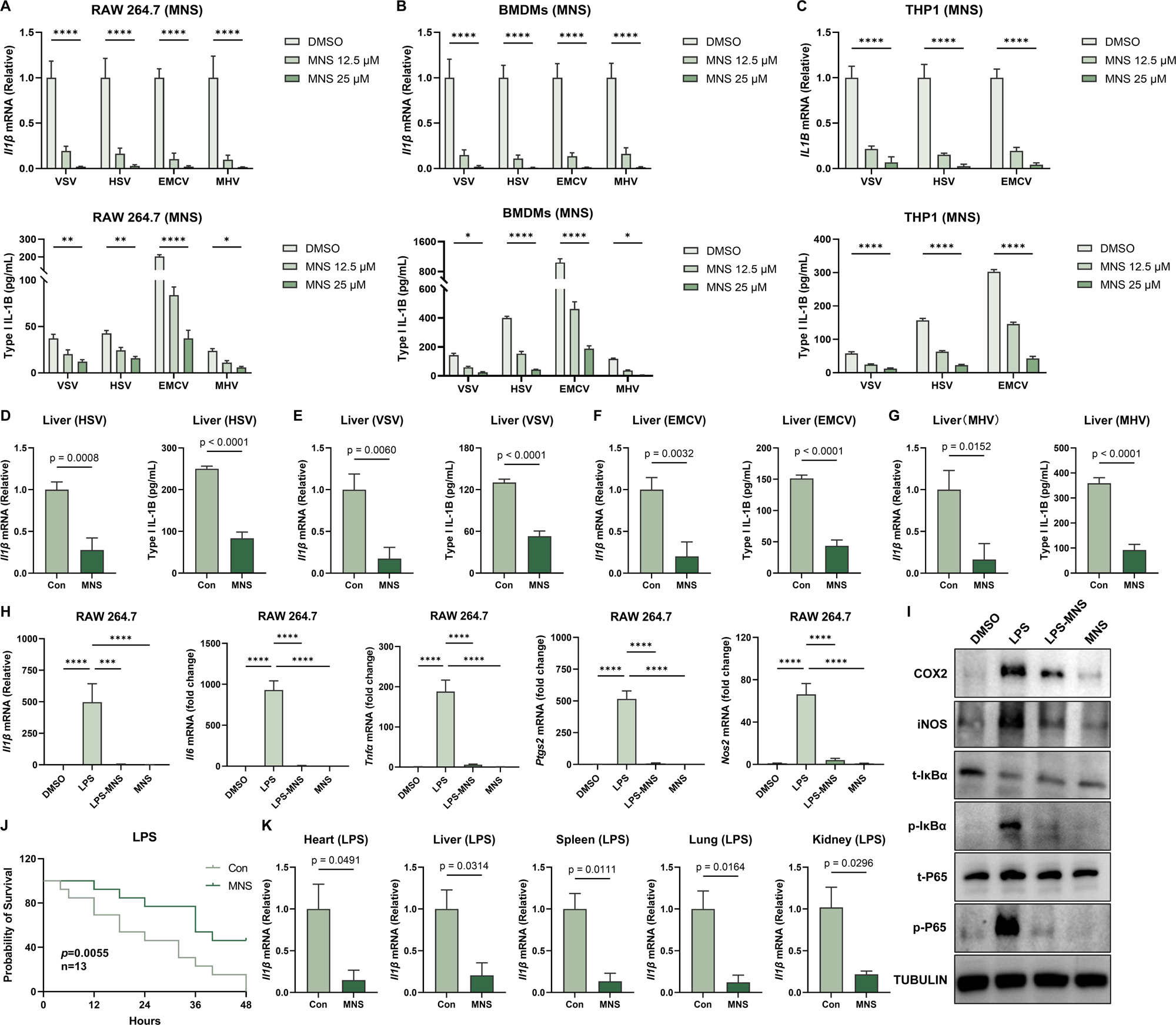
MNS Suppressed Virus-induced and LPS-induced Inflammatory Responses. (A-C) IL1B mRNA (qPCR) and protein levels (ELISA) in RAW 264.7 cells (A), BMDMs (B), and THP1 cells (C) following treatment with DMSO or MNS and infection with indicated viruses. (D-G) Il1b expression (qPCR) and protein levels (ELISA) in liver tissues from C57BL/6J mice under viral challenge and MNS treatment. (H) MNS markedly downregulated the expression of *Il1b*, *Il6*, *Tnf*α, *Ptgs2*, *Nos2* mRNA in RAW264.7 cells. (I) MNS significantly reduced the expression of COX2, iNOS, p-IκBα, p-P65 at the protein level. (J) MNS treatment led to a significant reduction in mortality among LPS-challenged mice (n=13). (K) The expression level of Il1b in heart, spleen, lung, kidney, and liver was quantified by RT-qPCR. Values are presented as mean ± SEM. Significance was indicated as follows: N.S., p > 0.05; *p < 0.05; **p < 0.01; ***p < 0.001; ****p < 0.0001.

To determine whether these anti-inflammatory effects also occurred in vivo, we measured cytokine responses in liver tissues from mice infected with HSV-1, VSV, EMCV or MHV. Across all infection models, MNS treatment significantly reduced hepatic *Il1b* mRNA levels as well as IL-1B protein abundance compared with untreated controls (Figure 3D-3G). The consistent reduction in both transcriptional and protein markers indicated that MNS effectively dampened excessive inflammatory signaling during systemic viral infection.

Moreover, we tested whether it also suppressed LPS-induced inflammation. RT-qPCR analysis in RAW 264.7 macrophages showed that LPS stimulation robustly upregulated multiple classes of inflammatory genes compared with DMSO controls, including pro-inflammatory cytokines *Il1b*, *Il6*, and *Tnfα* mRNA, as well as inflammatory effector enzymes *Ptgs2* and *Nos2*. Co-treatment with MNS markedly reduced the LPS-induced expression of all these genes, while MNS alone did not increase their basal transcription levels (Figure 3H). Consistently, western blots showed that MNS co-treatment reduced LPS-induced phosphorylation of NF-κB p65 and IκBα, as well as COX-2 and iNOS protein levels, supportng MNS’s anti-inflammatory effect (Figure 3I). Encouraged by the in vitro findings, we next evaluated whether MNS could ameliorate systemic inflammation in vivo using an LPS-induced sepsis model. Six-week-old C57BL/6J mice were pretreated with a single intraperitoneal dose of MNS (5 mg/kg). One hour later, they were subjected to a lethal challenge of LPS (20 mg/kg, i.p.). While all mice in the vehicle control group succumbed within 48 hours, approximately 50% of the animals receiving MNS survived (Figure 3J). To evaluate the systemic inflammatory response further, blood along with heart, liver, spleen, lung, and kidney tissues were collected 12 hours after LPS administration. MNS-treated mice exhibited significantly decreased levels of *Il1b* mRNA across all tested tissues compared to controls (Figure 3K). These results indicated that MNS effectively protected mice from hyperinflammation and endotoxic lethality, aligning with its potent anti-inflammatory action observed in vitro.

### Integrated Omics Analysis of MNS-Mediated Inflammation Suppression

To comprehensively evaluate how MNS modulated LPS-driven transcriptional responses in RAW264.7 macrophages, we performed bulk RNA-seq on all treatment groups. Heatmap visualization of differentially expressed genes (DEGs) showed that the transcriptional profile of the LPS+MNS group was clearly separated from LPS alone and displayed broadly reduced expression of inflammatory genes (Figure 4A). Consistent with this pattern, volcano plot analysis identified 2218 genes upregulated and 3364 genes downregulated by MNS co-treatment compared with LPS (Figure 4B). GO and KEGG enrichment revealed that these DEGs mainly involved inflammation-related pathways, including cytokine signaling, NF-κB activation, and innate immune responses (Figure 4C-4D).

**Figure 4.**
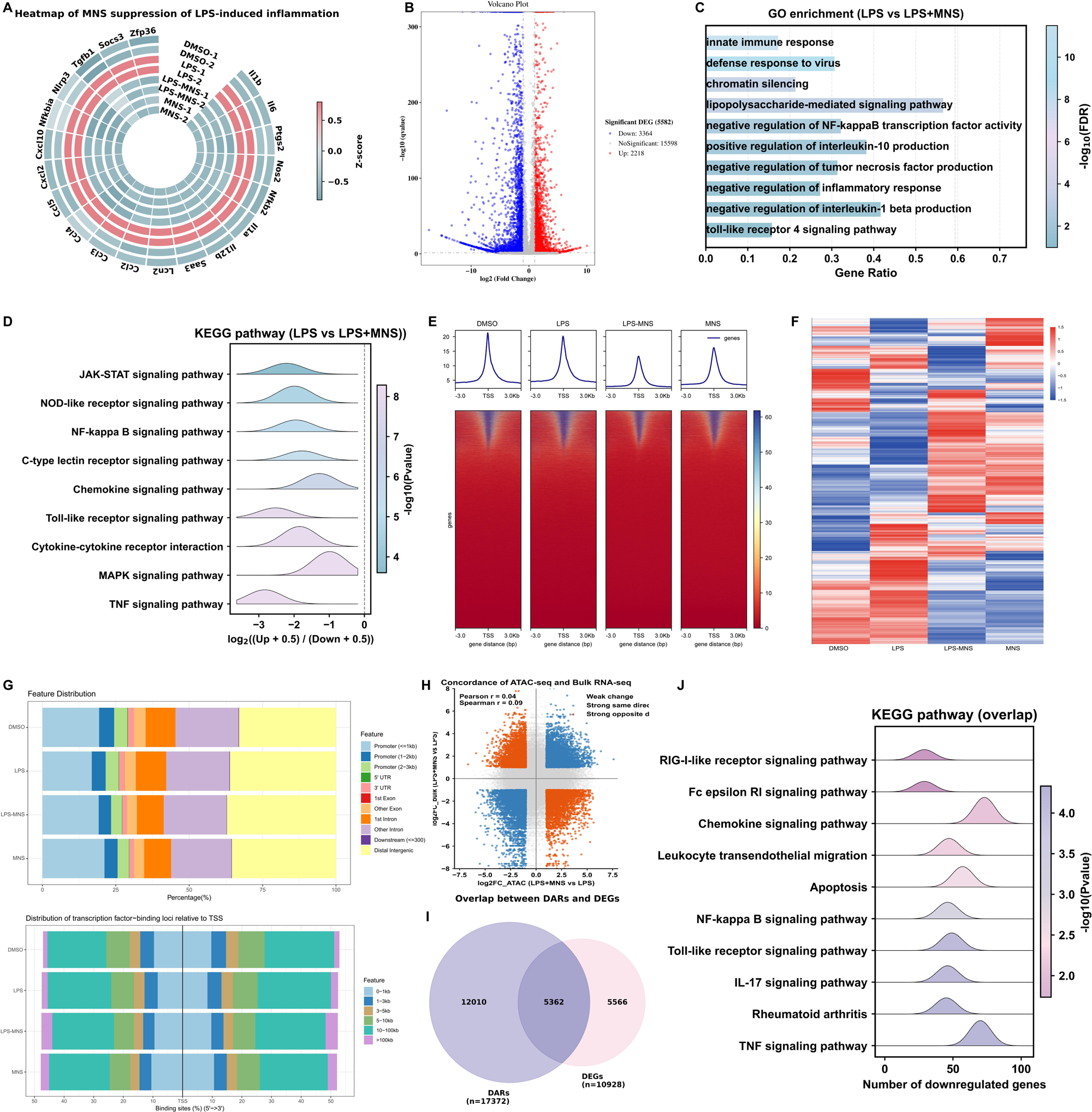
Integrated Omics Analysis of MNS-Mediated Inflammation Suppression. (A) Heatmap of inflammatory pathway gene expression in cells treated with DMSO, LPS, LPS+MNS, or MNS alone, highlighting MNS-mediated regulation relative to LPS. (B) Volcano plot showed DEGs between the treatment of LPS+MNS and LPS alone in RAW264.7 cells. (C) Gene Ontology (GO) enrichment of DEGs in (B). (D) KEGG enrichment of DEGs in (B). (E) Heatmap showed that MNS markedly changed the chromatin accessibility induced by LPS. (F) Heatmap illustrated the differentially accessible regions (DARs), following treatment with DMSO, LPS, LPS-MNS and MNS alone. (G) Distribution of peaks and transcription factor-binding loci relative to transcription start sites (TSS) in the LPS, LPS-MNS, and MNS groups compared to the DMSO group, respectively, as annotated using the ChIPseeker R package. (H) Scatter plot analysis comparing fold changes from ATAC-seq and RNA-seq. (I) Overlap downregulated genes in ATAC-seq and bulk RNA-seq between LPS and LPS-MNS groups in RAW 264.7 cells. (J) KEGG enrichment of overlap genes in (I).

To investigate the epigenetic basis underlying these transcriptional changes, we next performed ATAC-seq under DMSO, LPS, LPS+MNS, and MNS-alone conditions. LPS stimulation markedly increased global chromatin accessibility, with strong enrichment around transcription start sites (TSSs). This was evident from both the aggregate TSS profiles and the TSS-centered heatmaps, which showed widespread accessibility gains across promoters and gene bodies in the LPS group. Co-treatment with MNS clearly attenuated these LPS-induced accessibility increases, as reflected by reduced TSS signal intensity and visibly narrower accessible regions in the heatmaps. MNS alone did not cause global chromatin opening, indicating that its primary effect was the suppression of LPS-driven epigenomic activation (Figure 4E-4F).

We further examined the genomic distribution of accessible peaks. LPS increased accessibility across promoter regions, 5′UTRs, exons, introns, and distal regulatory regions, whereas MNS co-treatment shifted this distribution toward a pattern resembling the basal DMSO state. In addition, LPS enhanced the accumulation of transcription factor-binding sites near TSSs, while MNS reduced this enrichment and restored a distribution similar to resting macrophages (Figure 4G).

Finally, we integrated the RNA-seq and ATAC-seq datasets to identify coordinated transcriptional and chromatin changes. Scatter plot analysis comparing fold changes from ATAC-seq and RNA-seq revealed a weak but detectable overall relationship between chromatin accessibility and gene expression (Figure 4H). Intersection analysis showed that 10928 downregulated genes overlapped with 17372 loci exhibiting reduced promoter accessibility in the LPS+MNS group (Figure 4I). Enrichment analysis based on the KEGG pathway database demonstrated that these overlapping genes were markedly downregulated in several core inflammatory pathways, such as NF-κB, Toll-like receptor, TNF, and chemokine signaling (Figure 4J), indicating that MNS repressed inflammation through coordinated regulation at both the chromatin and transcriptional levels.

Together, these integrated multi-omics results demonstrated that MNS exerted potent anti-inflammatory activity by broadly suppressing LPS-induced gene expression and restraining the accompanying chromatin accessibility changes across key inflammatory pathways.

## Discussion

In this study, we identified the small-molecule nitrovinyl benzodioxole MNS as a host-directed agent that combined broad-spectrum antiviral activity with potent anti-inflammatory effects. Using multiple cell lines and four distinct RNA and DNA viruses, we showed that MNS reduced viral infection and viral gene expression and established a host-protective antiviral state in vitro. Consistent with these findings, MNS improved survival in several lethal viral infection models in mice, lowered systemic and tissue viral loads, and attenuated virus-induced tissue damage. In parallel, MNS dampened virus-induced and LPS-induced inflammatory responses in macrophages and multiple organs and protected mice from LPS-induced endotoxic lethality. Integrated bulk RNA-seq and ATAC-seq analyses further demonstrated that MNS broadly repressed LPS-driven inflammatory gene programs and reversed chromatin accessibility gains at promoters and transcription start sites of key inflammatory pathways. Together, these data supported MNS as a host-directed compound that simultaneously enhances antiviral defense and constrains hyperinflammatory responses.

A major unmet need in infectious and inflammatory diseases is the development of therapies that move beyond single-virus targets and address both viral burden and host-driven pathology. Most approved antivirals directly target viral enzymes or structural proteins, which limits their spectrum and makes them vulnerable to resistance when viruses accumulate escape mutations [41, 42]. Host-directed antiviral strategies seek instead to modulate cellular processes required for viral replication or to strengthen endogenous defense mechanisms. Several host-directed candidates are now being tested clinically, but most have been optimized either for antiviral efficacy or for anti-inflammatory effects, rather than for a balanced dual action. Our work addresses this gap by providing experimental evidence that a single small molecule can confer broad antiviral protection while at the same time limiting excessive inflammation in both virus-infected and endotoxemic settings.

Macrophages are central to this dual role. Upon detecting viral or bacterial components via pattern-recognition receptors, these cells promptly activate NF-κB and interferon signaling, leading to the production of cytokines and chemokines [43]. These responses are essential for pathogen control, but they can also drive systemic inflammation, vascular injury, and organ failure when sustained or dysregulated [44]. Recent reviews have emphasized the importance of balancing host defense and viral tolerance to prevent tissue damage during infection. Our data showed that MNS shifted this balance toward a more protective state, it suppressed infection of multiple viruses in macrophages and in vivo, yet concurrently lowered Il1b and other inflammatory mediators in infected tissues and improved survival in a severe sepsis model. This dual phenotype distinguishes MNS from many immune-stimulatory agents that primarily boost antiviral responses but may exacerbate cytokine-driven damage.

The broad-spectrum antiviral activity of MNS suggests a host-directed mechanism rather than virus-specific inhibition. Unlike direct-acting antivirals that target viral enzymes or structural proteins, MNS effectively restricts both RNA and DNA viruses across multiple experimental systems, indicating modulation of shared host antiviral pathways. Notably, MNS treatment resulted in a disproportionate reduction in infectious virus production relative to viral RNA abundance, pointing to interference with post-transcriptional steps of the viral life cycle. Such effects are consistent with interferon-stimulated host restriction mechanisms that impair viral protein synthesis, assembly, or virion maturation without immediately eliminating viral transcripts. In parallel, transcriptomic and chromatin accessibility analyses demonstrate that MNS reprograms innate immune responses by dampening excessive inflammatory signaling while maintaining antiviral defense programs. Together, these findings support a model in which MNS establishes a balanced, host-protective antiviral state that limits viral replication and pathology while avoiding immune-mediated tissue damage.

Mechanistically, MNS has been reported to function as an orally available tyrosine kinase inhibitor with broad-spectrum antiplatelet effects, targeting Src, Syk, and FAK while also suppressing NLRP3 inflammasome activation and β1 integrin signaling. In preclinical studies, various small-molecule NLRP3 inhibitors and other inflammasome-targeting agents have demonstrated therapeutic potential in models of ischemia–reperfusion injury, metaflammation, and similar inflammatory disorders [45, 46]. However, most of these molecules have been developed primarily as anti-inflammatory agents, and their antiviral properties have not been systematically examined. Our study filled this gap by demonstrating that a compound with known NLRP3- and kinase-inhibitory activity also exerted broad antiviral effects in vitro and in vivo. While we did not dissect each target individually, the convergence on Src/Syk/FAK signaling, NF-κB activation, and NLRP3 inflammasome function provided a plausible mechanistic framework for the combined antiviral and anti-inflammatory actions of MNS.

At the transcriptional and epigenetic levels, multiple studies have shown that macrophage activation is governed by dynamic remodeling of chromatin accessibility and transcription factor binding, which together shape inflammatory gene expression, tolerance, and trained immunity [47]. LPS and other inflammatory stimuli open chromatin at promoters and enhancers of NF-κB and interferon target genes, and the resulting epigenomic state can either amplify or restrain subsequent responses [48]. In this context, a key knowledge gap has been how small-molecule drugs with host-directed activity sculpt these chromatin landscapes in primary innate immune cells. By integrating RNA-seq and ATAC-seq, we showed that MNS not only downregulated inflammatory transcripts but also reduced LPS-induced chromatin accessibility around transcription start sites and promoters of genes involved in NF-κB, Toll-like receptor, and TNF signaling. This coordinated transcriptional and epigenomic repression suggests that MNS does not simply block one downstream effector, but instead resets the inflammatory gene network at the level of chromatin architecture.

Our integrated analysis further indicated a limited overall association between alterations in chromatin accessibility and corresponding gene expression changes, with a substantial number of genes exhibiting non-aligned or complex regulatory dynamics. This is consistent with recent work indicating that inflammatory gene regulation involves complex interactions between transcription factors, chromatin remodelers, and three-dimensional genome organization, and that accessibility changes do not always translate directly into steady-state mRNA levels [49]. Within this broader landscape, we found that genes showing both reduced promoter accessibility and downregulated expression were strongly enriched for core inflammatory pathways. These findings suggest that MNS preferentially converged on a subset of regulatory nodes where chromatin opening and transcription are tightly coupled, providing a focused yet system-level mechanism to dampen hyperinflammatory signaling.

Our sepsis data further support the concept that epigenetically mediated reprogramming of macrophage and tissue responses can translate into improved systemic outcomes. Protection from LPS-induced lethality, along with reduced Il1b expression across multiple organs, aligns with recent evidence that modulating NLRP3 inflammasome activity and NF-κB signaling can ameliorate organ injury in models of endotoxemia and ischemia–reperfusion [50, 51]. Compared with selective NLRP3 inhibitors that mainly target inflammasome assembly, MNS appears to act at several levels of the inflammatory cascade, including upstream kinase pathways, NF-κB activation, and chromatin accessibility at inflammatory loci. This broader mechanism may offer advantages in settings where multiple parallel pathways drive hyperinflammation, although it may also carry distinct safety considerations.

From a translational perspective, the combined antiviral and anti-inflammatory profile of MNS suggests potential applications in viral infections characterized by strong inflammatory components, such as severe influenza, viral sepsis, and other acute viral pneumonias. Host-directed agents are particularly attractive for newly emerging pathogens, where direct-acting antivirals may not yet exist and rapid viral evolution can undermine target-specific drugs [18]. By targeting host pathways that are shared across viruses, and by preventing the host response from overshooting, MNS-like compounds could help bridge the gap between early outbreak management and later development of pathogen-specific therapies.

This study also has several limitations that point to important directions for future work. First, although we used multiple viruses and cell types, our panel did not include clinically important pathogens such as influenza, coronaviruses, or highly pathogenic emerging viruses. Testing MNS in additional viral models, including those with distinct tissue tropism and immune evasion strategies, will be necessary to define the true breadth of its antiviral spectrum. Second, we inferred mechanisms largely from pathway-level readouts, kinase targets reported in prior work, and genome-wide expression and accessibility profiles. Direct target-engagement studies, phosphoproteomic analyses, and genetic perturbation of candidate pathways, such as Src/Syk/FAK or NLRP3, will be needed to clarify which molecular interactions are essential for the observed phenotypes. Third, MNS has been reported to possess antiplatelet activity and cytotoxicity at certain concentrations. A detailed assessment of its therapeutic window, off-target effects, and pharmacokinetics in models of viral infection and sepsis will be required before clinical translation can be considered.

In addition, our multi-omics analysis focused on bulk macrophage populations and whole tissues. Single-cell RNA-seq and single-cell ATAC-seq could provide finer resolution of how MNS affects distinct immune and stromal cell subsets, and whether it preferentially reprograms specific macrophage states, such as inflammatory versus tissue-repair phenotypes. Integrating these approaches with spatial transcriptomics or imaging-based techniques would help to map where in tissues MNS exerts its dominant effects and how this relates to local viral replication and tissue damage.

Despite these limitations, our findings expand the current landscape of host-directed therapies by demonstrating that MNS can couple broad antiviral activity with epigenetically mediated suppression of hyperinflammatory responses. By linking functional virology and sepsis models with integrated RNA-seq and ATAC-seq, we begin to fill an important gap in understanding how small molecules can reshape inflammatory gene networks at both transcriptional and chromatin levels. Further optimization of MNS or related scaffolds, together with more detailed mechanistic and safety studies, may ultimately yield new therapeutic options for viral infections and hyperinflammatory diseases where both pathogen control and immune restraint are required.

## Materials and methods

### Reagents and antibodies

Rabbit anti-COX2 antibody was obtained from Wanleibio. Primary antibodies from Proteintech comprised rabbit anti-iNOS (catalog 80517-1-RR), rabbit anti-β-Tubulin (catalog 10094-1-AP), and mouse anti-GFP tag (catalog 50430-2-AP). Additional antibodies were purchased from Cell Signaling Technology, including rabbit anti-phospho-NF-κB p65 (Ser536) (clone 93H1; catalog 3033), anti-NF-κB p65 (D14E12) XP (catalog 8242), anti-phospho-IκBα (Ser32) (clone 14D4; catalog 2859), and anti-IκBα (clone 44D4; catalog 4812). Secondary antibodies consisted of HRP-conjugated Affinipure Goat Anti-Rabbit IgG (H+L) (catalog SA00001-2) and HRP-conjugated Affinipure Goat Anti-Mouse IgG (H+L) (catalog SA00001-1), both from Proteintech. Dimethyl sulfoxide (DMSO; catalog D8371) was purchased from Solarbio. MNS (catalog HY-78263) and recombinant mouse GM-CSF (catalog HY-P7361) were obtained from MedChemExpress.

### Cells

All cell lines (RAW264.7, THP1, HT1080, 17Cl-1, Vero) were acquired from the American Type Culture Collection (ATCC). To generate BMDMs, bone marrow was harvested from femurs and tibiae of 8-week-old C57BL/6J mice and then differentiated with GM-CSF for 7 days. Both the acquired cell lines and the derived BMDMs were cultured in Dulbecco’s Modified Eagle’s Medium (DMEM) (03.1002C, EallBio). The medium was consistently supplemented with 10% fetal bovine serum (FBS) and 1% penicillin-streptomycin to support normal growth and maintenance.

### Virus infection and propagation

Cells were infected at 70–80% confluence with the indicated multiplicities of infection (MOI): vesicular stomatitis virus (VSV, Indiana strain, MOI 0.1), herpes simplex virus type 1 (HSV-1, F strain, MOI 0.5), encephalomyocarditis virus (EMCV, MOI 0.1), and mouse hepatitis virus (MHV, A59 strain, MOI 0.1). In addition, GFP-labelled VSV (MOI 0.1), HSV (MOI 0.1) and vaccinia virus (VACV, MOI 0.1) were used.

Notably, consistent with the high permissiveness of RAW 264.7 macrophages and the rapid multi-cycle replication kinetics of VSV, high viral titers were observed at early time points in control cultures.

For preparation of virus stocks, VSV Indiana and VSV-GFP (kindly provided by J. Rose, Yale University) were propagated in Vero cells. HSV-1 strain 17, VSV-GFP, HSV-GFP and VACV-GFP (from Zhengfan Jiang, Peking University) were also expanded in Vero cells. MHV-A59 (ATCC VR-764) and EMCV (ATCC VR-129B) were maintained in 17Cl-1 and Vero cells, respectively.

### Mice and in vivo virus infection

All animal experiments were conducted in compliance with the guidelines issued by the Chinese Association for Laboratory Animal Science and received approval from the Animal Care and Use Committee of Peking University Health Science Center (Ethics Approval Number: LA2016240). Every effort was taken to reduce animal discomfort and to employ the fewest animals necessary for statistically valid results.

All experiments utilized wild-type C57BL/6J mice sourced from the Department of Laboratory Animal Science, Peking University Health Science Center. These mice were maintained under specific pathogen-free conditions within the university’s Laboratory Animal Center. For the in vivo studies, age- and sex-matched littermates between 6 and 8 weeks old were utilized. Mice were intraperitoneally injected with various viruses at a dose of 1 × 10□ PFU per mouse, and survival was recorded daily.

### Euthanasia and anesthesia

Mice were anesthetized with isoflurane prior to tissue collection. In compliance with the AVMA Guidelines for the Euthanasia of Animals (2020), animals were euthanized at experimental endpoints using CO□ inhalation followed by cervical dislocation. All deaths occurred within the planned humane endpoints, with no unexpected fatalities.

### Construction of a sepsis model via LPS injection

Sepsis induced by lipopolysaccharide (LPS) was used to model systemic inflammatory responses. Male C57BL/6J mice aged 6 to 8 weeks and weighing 20 to 25 grams were randomly assigned to two groups with twelve animals per group, an LPS plus PBS group and an LPS plus MNS group. After i.p. injection of LPS (20 mg/kg), mice were administered MNS (5 mg/kg, i.p.) or PBS. Survival was monitored over 48 h, with blood and tissues harvested 12 h post-LPS.

### Hematoxylin-eosin (H&E) staining

Collected tissues underwent cold□saline rinsing, 4% paraformaldehyde fixation, dehydration, paraffin embedding, and 5□μm sectioning before being stained with hematoxylin and eosin (H&E).

### RNA extraction, reverse transcription and quantitative PCR

Total RNA was isolated from stimulated or virus-infected cells using TRIzol reagent (TIANGEN, A0123A01) following the supplier’s protocol. In brief, cells were lysed in TRIzol, homogenized and subjected to phase separation; the aqueous phase was collected, and RNA was precipitated, washed and finally dissolved in RNase-free water. RNA concentration and purity were assessed spectrophotometrically (A260/A280). For reverse transcription, 0.5–1 μg RNA was converted to cDNA using HiScript II RT SuperMix (Vazyme, R223-01). Quantitative PCR was performed in 96-well plates with SYBR Green qMix (Vazyme, Q311). Gene expression levels were normalized to Actb and calculated using the 2^(-ΔΔCt) method. Primer sequences are listed in Supplementary Table S1.

### Total protein extraction and western blotting

Cells were lysed in strong RIPA buffer (MedChemExpress, HY-K1001) supplemented with an EDTA-free protease inhibitor mix and phosphatase inhibitor cocktails II and III (100×). Lysates were cleared by centrifugation, and 10–30 μg of total protein per sample was separated by SDS–PAGE and electrophoretically transferred to nitrocellulose membranes (Beyotime, FFN08). Membranes were blocked in TBST containing 5% skim milk or 5% BSA, incubated with the appropriate primary antibodies, and then with HRP-conjugated secondary antibodies. Protein bands were visualized using enhanced chemiluminescence (ECL; EallBio, 07.10009-50).

### Enzyme-linked immunosorbent assay (ELISA)

IL1B levels in cell culture were measured from the supernatants of RAW264.7 cells, BMDMs and THP-1 cells after MNS stimulation. Cell culture supernatants were collected, centrifuged at 1600 rpm for 5 minutes to remove debris, and stored at -80°C until analysis. Mouse cells were assayed with a mouse IL1B ELISA kit (Boster, EK2285), and human primary cells were assayed with a human IL1B ELISA kit (Boster, EK2286), following the manufacturers’ instructions. Absorbance was read at 450 nm on a microplate reader, and cytokine concentrations were calculated from the standard curve.

For tissue cytokine measurement, tissues were collected at the indicated time points, rinsed briefly in cold PBS to remove residual blood, and weighed. Tissues were homogenized in cold PBS supplemented with protease inhibitors using a handheld homogenizer, then centrifuged at 12000 rpm for 10 min at 4°C. The clarified supernatants were used for ELISA. IL1B in tissue homogenates was quantified with the same mouse IL1B ELISA kit, according to the manufacturers’ protocols. When indicated, cytokine levels were normalized to tissue weight or total protein content to allow comparison between animals.

### Plaque assay

Vero cells were grown to confluence in 24-well plates and infected the following day. For quantifying virus in culture supernatants, samples were harvested at the specified time points, clarified by centrifugation (1600 rpm, 5 min) and then serially diluted 10-fold in DMEM. Tissue specimens were weighed, homogenized in chilled DMEM or PBS, centrifuged at 3000 rpm for 10 min at 4 °C to remove debris, and the resulting supernatants were likewise subjected to 10-fold serial dilutions.

For plaque formation, 200 μL of each virus dilution was added to Vero monolayers and incubated for 1 h at 37 °C with gentle mixing every 15 min. After adsorption, inocula were removed, cells were rinsed with PBS, and overlaid with DMEM containing 0.5% CMC (Sigma, 419338) and 2% FBS. Following a 48-h incubation at 37 °C, cells were fixed in 4% paraformaldehyde for 20 min and stained with 1% crystal violet to visualize plaques. Viral titers (PFU/mL) were calculated from plaque numbers, the corresponding dilution factor, and the volume of inoculum.

### Fluorescence assay

Cells were infected with GFP-tagged viruses at an MOI of 0.1 for 12 hours in the presence or absence of MNS. After incubation, cells were gently washed with PBS to remove unattached virus and were kept in fresh complete medium. Images of GFP signals were acquired at 250× magnification on a Nikon fluorescence microscope (excitation 488□nm, emission 510□nm) under uniform exposure conditions, allowing reliable intergroup comparison.

### RNA-seq and data analysis

Total RNA was extracted with a high□throughput kit (YEASEN, 19211ES60). Subsequent library preparation, sequencing, and primary analysis followed the protocols of Suzhou GENEWIZ Biotechnology Co., Ltd. featureCounts (v2.0.0) was used to generate gene counts, which were normalized to FPKM. Differential expression was assessed via DESeq2 (R v1.38.3) with cut□offs of |log□(fold change)| > 2 and adjusted p < 0.05.

### ATAC-seq

ATAC-seq was performed to profile genome-wide chromatin accessibility using a transposase-based workflow. Briefly, nuclei were isolated from cells and incubated with Tn5 transposase preloaded with sequencing adapters, allowing preferential tagging of open chromatin regions. Adapter-tagged fragments were then PCR-amplified with indexed primers to generate sequencing libraries. Libraries were prepared with the Hyperactive ATAC-Seq Library Prep Kit (TD711, Vazyme Biotech) and sequenced by GENEWIZ Biotechnology (Suzhou, China). The resulting datasets were used to identify regions of accessible chromatin across the genome.

### Data analysis of ATAC-seq

For ATAC-seq high-throughput data generated by GENEWIZ (Suzhou, China), raw FASTQ files were first subjected to quality control and adapter trimming with Trim Galore (v0.6.4). Clean reads were aligned to the mouse genome (mm10) using bowtie2 (v2.3.5.1) with a very sensitive setting and an insert-size limit of 2,000 bp. Peaks were called for each sample with MACS3 (v3.0.0a5) under default parameters to obtain BED-formatted peak files. To generate a unified set of accessible regions across samples, individual ATAC-seq peak files were merged with bedtools merge (v2.31.1). Signal tracks were produced and normalized with deepTools bamCoverage (v3.3.2) using the options --binSize 100 --normalizeUsing RPKM --effectiveGenomeSize 2864785220 --ignoreForNormalization chrM --extendReads, i.e. bamCoverage --bam $var -o ${var%.*}.bw --binSize 100 --normalizeUsing RPKM --effectiveGenomeSize 2864785220 --ignoreForNormalization chrM –extendReads [52]. Differential accessibility between treatment and control groups was assessed in R with the csaw package (v1.38.0), and peak annotation to nearby genes and regulatory features was performed with ChIPseeker (v1.34.1). Final bigWig tracks and representative loci were inspected in Integrative Genomics Viewer (IGV, v2.17.4) to confirm the ATAC-seq peak patterns.

### Quantification and statistical analysis

Analyses used GraphPad Prism. Two□group comparisons relied on unpaired two□tailed t□tests (brackets in figures). Multi□group/multi□factor comparisons used one□ or two□way ANOVA plus Tukey’s post□hoc test. Survival was assessed by log□rank (Mantel–Cox) test. Bar graphs showed one of ≥3 independent experiments. Data were mean ± SEM (n ≥ 3). Significance: n.s., p > 0.05; *p < 0.05; **p < 0.01; ***p < 0.001; ****p < 0.0001.

## Data Availability Statement

The data that support the findings of this study are available from the corresponding author upon reasonable request. All sequencing data have been deposited in Sequence Read Archive under accession number PRJNA1366804.

## Acknowledgments

We thank the assistance of the Peking University High-Performance Computing Platform for providing access to computing clusters, which greatly facilitated the data analysis in this study.

## Author contributions

Y. Z. and F.Y. and H.Q. conceptualized, designed, and coordinated the study. Y.X. and X.C. and Y.Z. conducted the experiments. H.Q. and S.Z. and B.K. involved in data analysis and visualization. H.L. provided tools and reagents. X.C. modified the schematic diagram. Y.Z. and H.Q. wrote the manuscript, X.C. and W.Q. read and revised the final manuscript.

## Figure Legends

**Supplementary Figure 1.** (A) Dose-response curves showing the concentration-dependent inhibition of viral replication by MNS. Half-maximal effective concentrations (EC50) were calculated by nonlinear regression analysis. (B) Cell viability of RAW 264.7 cells and HT1080 cells treated with increasing concentrations of MNS in the absence of viral infection, assessed using a luminescence-based viability assay. Half-maximal cytotoxic concentrations (CC50) were determined by nonlinear regression. Data are mean□±□SEM; N.S., p□>□0.05; *p□<□0.05, **p□<□0.01, ***p□<□0.001.

**Supplementary Table 1. Primer sequences used for RT-qPCR.**

Forward and reverse primer sequences (5’→3’) for all genes analyzed in this study.

